## Supplementary Information for "Multi-parametric thrombus profiling microfluidics detects intensified biomechanical thrombogenesis associated with hypertension and aging"

Misbahud Din *et al.*

**This file includes:**

Supplementary Methods

Supplementary Figures: 1 to 12

Supplementary Tables: 1 to 3

**Supplementary Methods**

*Fluid dynamics simulation and calculation*

Blood as fluid was modeled using the inelastic non-Newtonian Carreau-Yasuda model^1^, which is commonly used in hemodynamical simulations^1-3^. In this model, the apparent viscosity is expressed as

$\mu\left( \dot{\gamma} \right)=\mu_{\infty}+\left( \mu_{0}-\mu_{\infty} \right){[1+{(\lambda\dot{\gamma})}^{\alpha}]}^{{(n-1)}/\alpha}$	(1)

in which $\mu_{0}=0.1600 Pa\cdot s$ is the viscosity under zero shear rate, $\mu_{\infty}=0.0035 Pa\cdot s$ is the viscosity under infinite shear rate, and model parameters $\lambda=8.2 s$, n =0.2128, and $\alpha=0.64$ are empirically determined constants. Values of the above parameters were all adopted from Ref^3^.

Fluid simulation was performed in three tubes with the same length (500 µm) but different cross-sectional geometries: rectangle (200 µm × 50 µm, identical to that of the devices used in experiments), square (100 µm × 100 µm), and circle (R = 56.42 µm), which have the same cross-section area. All three tubes contain a site of stenosis of the same level (S = 80%) and a similar shape (identical contraction angle and narrowing length). To mitigate convergence challenges in the vicinity of sharp corners, manual filleting is employed to smooth out each of these sharp corners.

Software COMSOL Multiphysics 6.0 was utilized to computationally simulate the flow profile, wherein CFD module was used to solve the incompressible Navier Stokes (N-S) equation:

$\rho\left( \vec{u}\cdot\nabla\vec{u} \right)=-\nabla p+\mu\nabla^{2}\vec{u}$	(2)

where $\vec{u}$ is the velocity vector, $p$ is the pressure and $\rho=1060 kg/m^{3}$ is the fluid density. At the microchannel inlet and along the walls, a zero-gauge pressure boundary condition and a no-slip boundary condition were imposed, respectively. To ensure precision, an extra-fine CFD mesh consisting of approximately 8,000,000 elements was employed for each simulation.

Notably, the simulation also revealed an advantage of using the rectangular geometry for the channel over the square geometry used in some previous works, because the latter requires a lower perfusion rate to reach the same WSS (Supp. Fig. 1c,e,f), which can cause steplike movement of the syringe pump and undesired pulsatile flow.

The Reynolds number is calculated as $Re=\frac{\rho UL}{\mu}$, where $\rho$ is the fluid density, *U* is the average flow speed of the cross-section, and *L* is the characteristic length defined as $L= \frac{4A_{C}}{P}$, with $A_{C}$ and *P* representing the cross-sectional area and perimeter, respectively. Due to the high shear rate, the viscosity *μ* is approximated as $\mu_{\infty}$, the viscosity under infinite shear rate.

*Statistical analysis on data derived from clinical samples*

To compare thrombus profiling results from healthy young, healthy older, hypertensive young and hypertensive older adult groups, two-way ANOVA with unequal sizes and heteroscedastic factorial designs was used^4^. A Box-type finite-sample approximation method was applied for the distribution of quadratic forms^5^. Specifically, we denote the size of each group as $n_{\mathrm{ij}}$(i =1 (young) or 2 (older) and j =1 (healthy) or 2 (hypertensive)). Assume that the data points in each group are independent and follow the normal distribution $N \left( \mu_{ij}, \sigma_{ij}^{2} \right) (i=1,2, j=1,2)$, where $(\mu_{11},\mu_{12},\mu_{21},\mu_{22})$ are the respective means of different groups and $(\sigma_{11}^{2},\sigma_{12}^{2},\sigma_{21}^{2},\sigma_{22}^{2})$ are the respective variances of different groups. According to the model assumption of a two-way ANOVA:

$\mu_{ij}={\mu_{0}+\alpha}_{i}+\beta_{j}+\gamma_{ij} \left( i=1,2, j=1,2 \right)$	(3)

where $\mu_{0}$ is the common mean, $\alpha_{i}$ denotes the main effect of the hypertension factor, $\beta_{j}$ denotes the main effect of the aging factor, and $\gamma_{ij}$ denotes the interaction effect from both factors. The null hypotheses for two main effects and one interaction effect are as follows:

$$H_{01}:\alpha_{1}=\alpha_{2}=0$$

$$H_{02}:\beta_{1}=\beta_{2}=0$$

$$H_{03}:\gamma_{11}=\gamma_{12}=\gamma_{21}=\gamma_{22}=0$$

For each effect, the quadratic form statistics are calculated^5^. Let I be a $2\times2$ identity matrix and J a $2\times2$ matrix of 1’s. The symbol $\otimes$ denotes the Kronecker product of two matrices. We define $M_{1}=P\otimes\frac{1}{2}J$, $M_{2}=\frac{1}{2}J\otimes P$, and$M_{3}=P\otimes P$, where $P=I-\frac{1}{2}J$. Then, the above three null hypotheses of interest can be written using the unified form of

$$H_{0k}: M_{k}(\mu_{11},\mu_{12},\mu_{21},\mu_{22})'=0$$

for $k=1,2,3.$ The sample mean and sample covariance of data in each group are defined as $\bar{X}$ and $\hat{S}$, respectively, and it is further defined that matrix $D_{k}=diag(M_{k})$. Then the quadratic form test statistics for the k-th effect is defined as $F_{k}=\frac{N\cdot\bar{X}^{'}M_{k}\bar{X}}{tr(D_{k}\hat{S} )}$ for $k=1,2,3.$ According to the Box-type approximation^6^, the distribution of $F_{k}$ can be approximated by an F distribution $F_{(\hat{f}_{k},\hat{f}_{0k})}$, where $\hat{f}_{0k}=\frac{\left( \mathrm{tr}\left( D_{k}\hat{S} \right) \right)^{2}}{tr(D_{k}^{2}\hat{S}^{2}\Lambda)}$ and $\hat{f}_{k}=\frac{\left( \mathrm{tr}\left( D_{k}\hat{S} \right) \right)^{2}}{tr(M_{k}\hat{S}M_{k}\hat{S})}$, in which $\Lambda=diag\left\{ \frac{1}{n_{11}-1},\frac{1}{n_{12}-1},\frac{1}{n_{21}-1},\frac{1}{n_{22}-1} \right\}$.

**Supplementary Figures**

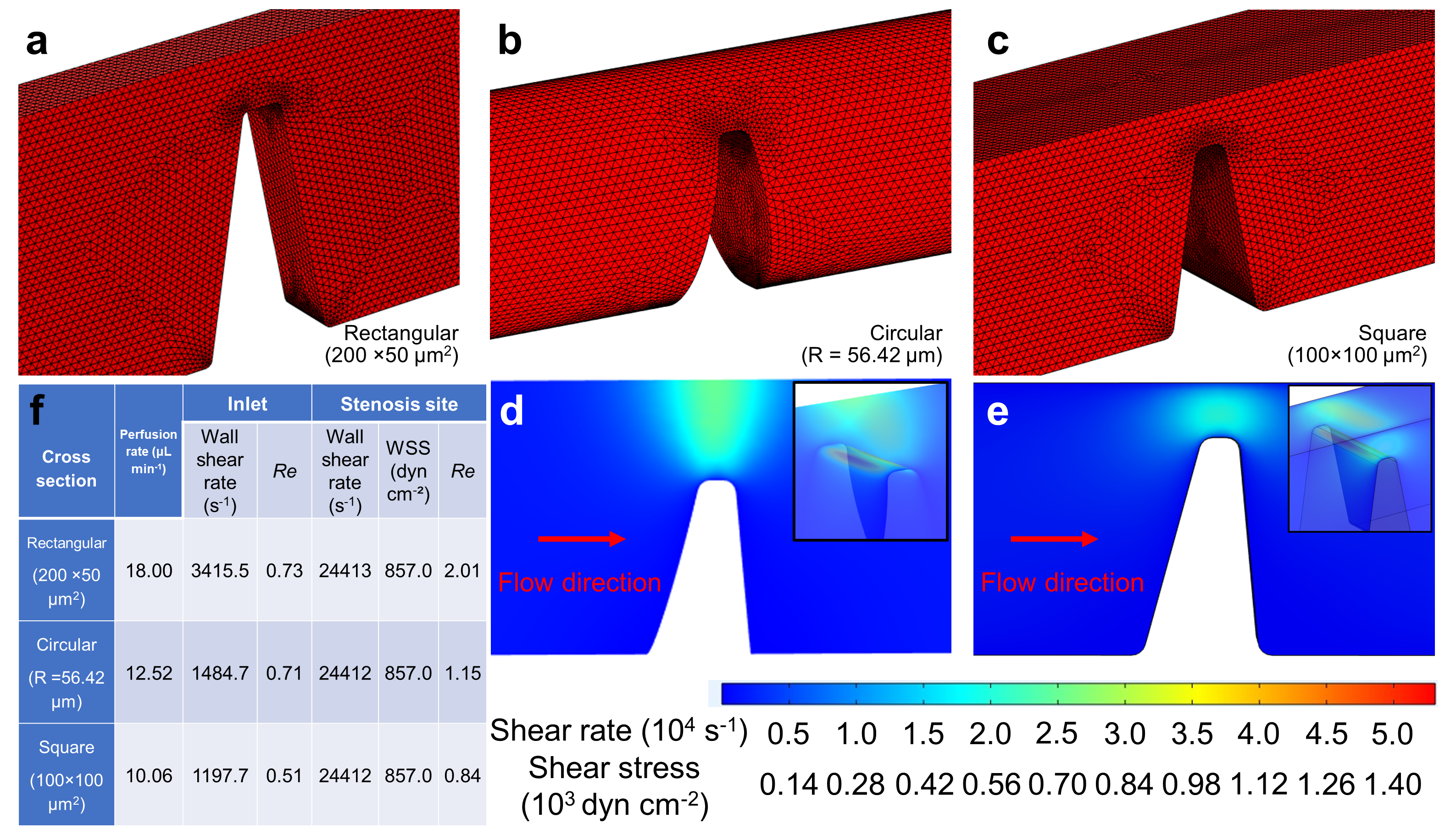

Supplemental Figure 1. Using fluid dynamics simulation to estimate the shear rate and shear stress distributions in stenotic channels with different designs. (a-c) Mesh plots of three designs of the stenotic channel, with the cross section being rectangular (a), circular (b) and square (c), respectively. In all three designs, the extent of stenosis is kept as 80%, and the angles of the narrowing and widening sides of the stenotic region are respectively 75° and 85°. (d,e) Simulated shear rate and shear stress distributions surrounding the stenosis area in the channel with round (d) and square (e) shaped designs, respectively. (f) Wall shear rate at the inlet, and wall shear rate and wall shear stress at the site of stenosis in the three stenotic channels when using the specified flow rates for blood perfusion.

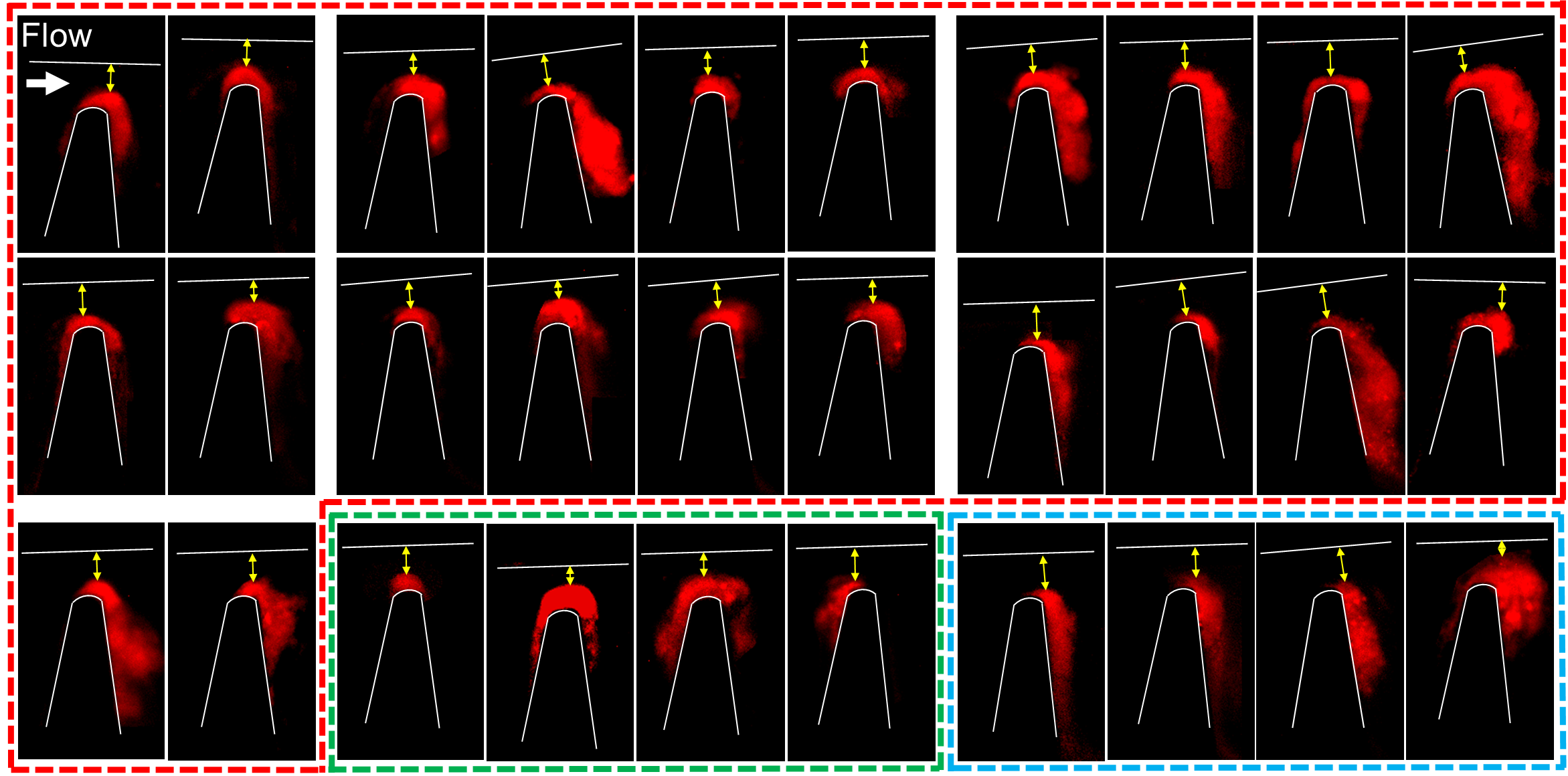

**Supplemental Figure 2.** **Representative** **pseudo-colored images of 30 healthy young subjects’ thrombi formed in the stenotic channels**. The margins of each channel are marked by white lines and curves. Among the thrombi, 26 out of 30 (87%) thrombi have a tendency of growing towards the downstream side of the stenosis (*surrounded by dashed red and blue lines*), which include 22 (73%) that cover the whole stenosis apex and 4 (13%) that do not, while the remaining 4 (13%) thrombi have the main body right on the apex or at the upstream side of the stenosis (*surrounded by dashed green lines*). A total of 26 (87%) thrombi have the point in their contour most close to the opposing channel wall (marked by yellow arrows) positioned above the apex of the stenosis (*surrounded by dashed red and green lines*), while 4 (13.3%) thrombi have their part most close to the opposing channel wall at the downstream side of the stenosis apex (*surrounded by dashed blue lines*). A total of 26 out of 30 (87%) thrombi cover the whole stenosis apex.

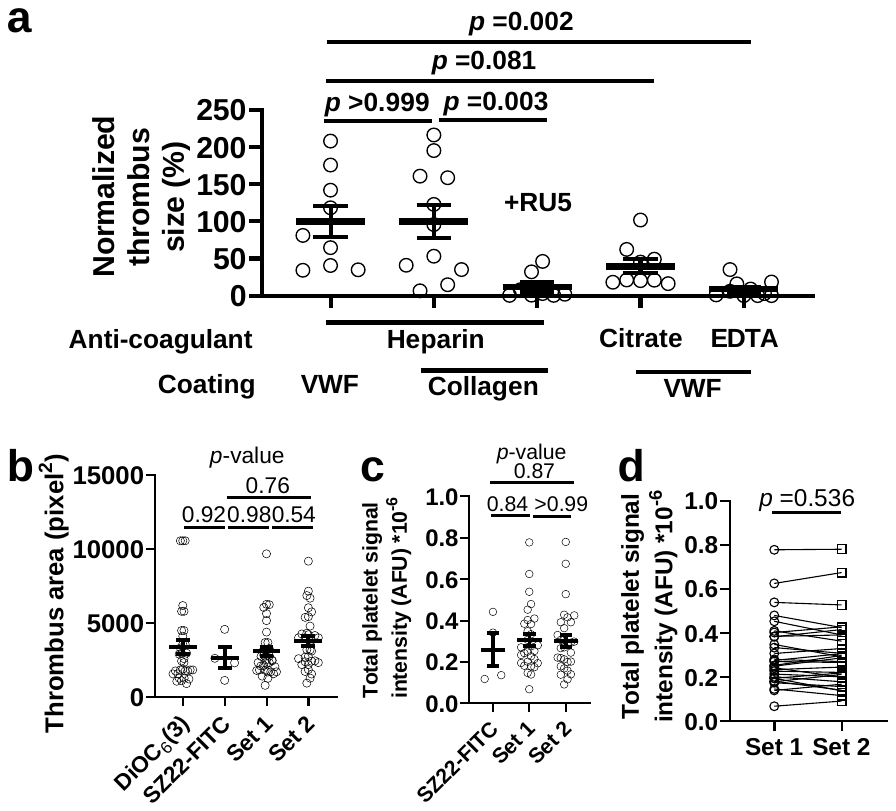

Supplemental Figure 3. Checking the impact of different anticoagulants, coating conditions and dyes on biomechanical thrombus formation. (a) Scatter plots with mean±s.e.m. of the normalized size of thrombi formed in blood anti-coagulated with heparin, citrate or EDTA, and in stenotic channels coated with VWF or collagen. In some runs with collagen coating, blood was added with 10 µg/ml RU5 to block plasma VWF binding to collagen. Data in all groups are normalized by the average of the heparin/VWF group. Each group contains data collected from at least 4 subjects. *P*-values: results of one-way ANOVA (F-value =8.391, degrees of freedom =46) and multiple comparison. (b) Scatter plots with mean±s.e.m. of the area of thrombi stained by DiOC_6_(3), SZ22-FITC, Sensor Set 1 and Sensor Set 2. *P*-values: results of one-way ANOVA (F-value =0.758, degrees of freedom =104) and multiple comparison. (c) Scatter plots with mean±s.e.m. of the size (total platelet signal intensity) of thrombi stained by SZ22-FITC, Sensor Set 1 and Sensor Set 2. *P*-values: results of one-way ANOVA (F-value =0.157, degrees of freedom =63) and multiple comparison. (d) Comparing the size of thrombi respectively stained by Sensor Set 1 and Sensor Set 2. Two points connected by a straight line denote two thrombi generated using the same blood sample. *P*-value: results of ratio paired t-test (*t*-value =0.626, degrees of freedom =29).

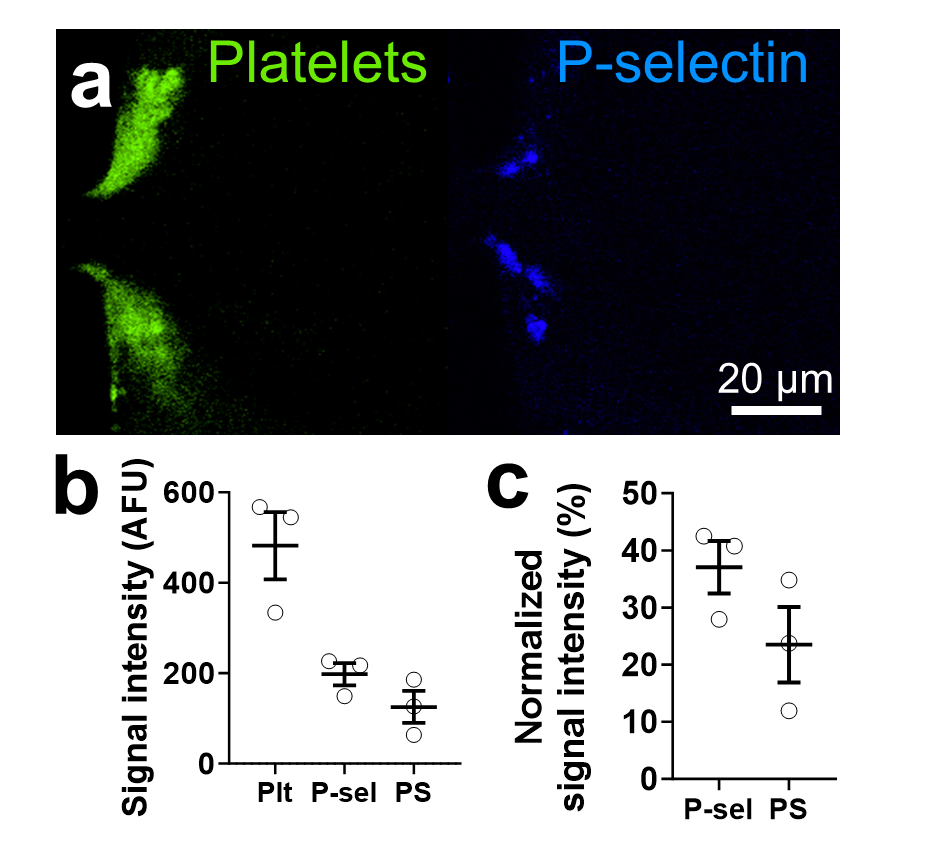

Supplemental Figure 4. Confirming the positive signals of P-selectin and PS in the biomechanical thrombi using a different stenosis channel design and microscope setup. Whole blood was added with either P2 (targeting integrin α_IIb_β_3_)-Alexa Fluor® 488 and AK4-Alexa Fluor® 647, or AK4-Alexa Fluor® 647 and Annexin V-Alexa Fluor® 488, and perfused through a double-hump channel with 80% stenosis. A confocal microscope with a motorized filter wheel was used for multi-fluorescence imaging. (a) Representative fluorescent images of thrombi stained with P2-Alexa Fluor® 488 targeting platelet (*green*) and AK4-Alexa Fluor® 647 targeting P-selectin (*blue*). (b) Scatter plots with mean±s.e.m. (n =3) of the signal intensities of biomarkers against platelets, P-selectin and PS 7.5 min after the onset of thrombus formation. (c) Scatter plots with mean±s.e.m. (n =3) of the signal intensities of P-selectin and PS normalized over platelet signal intensity.

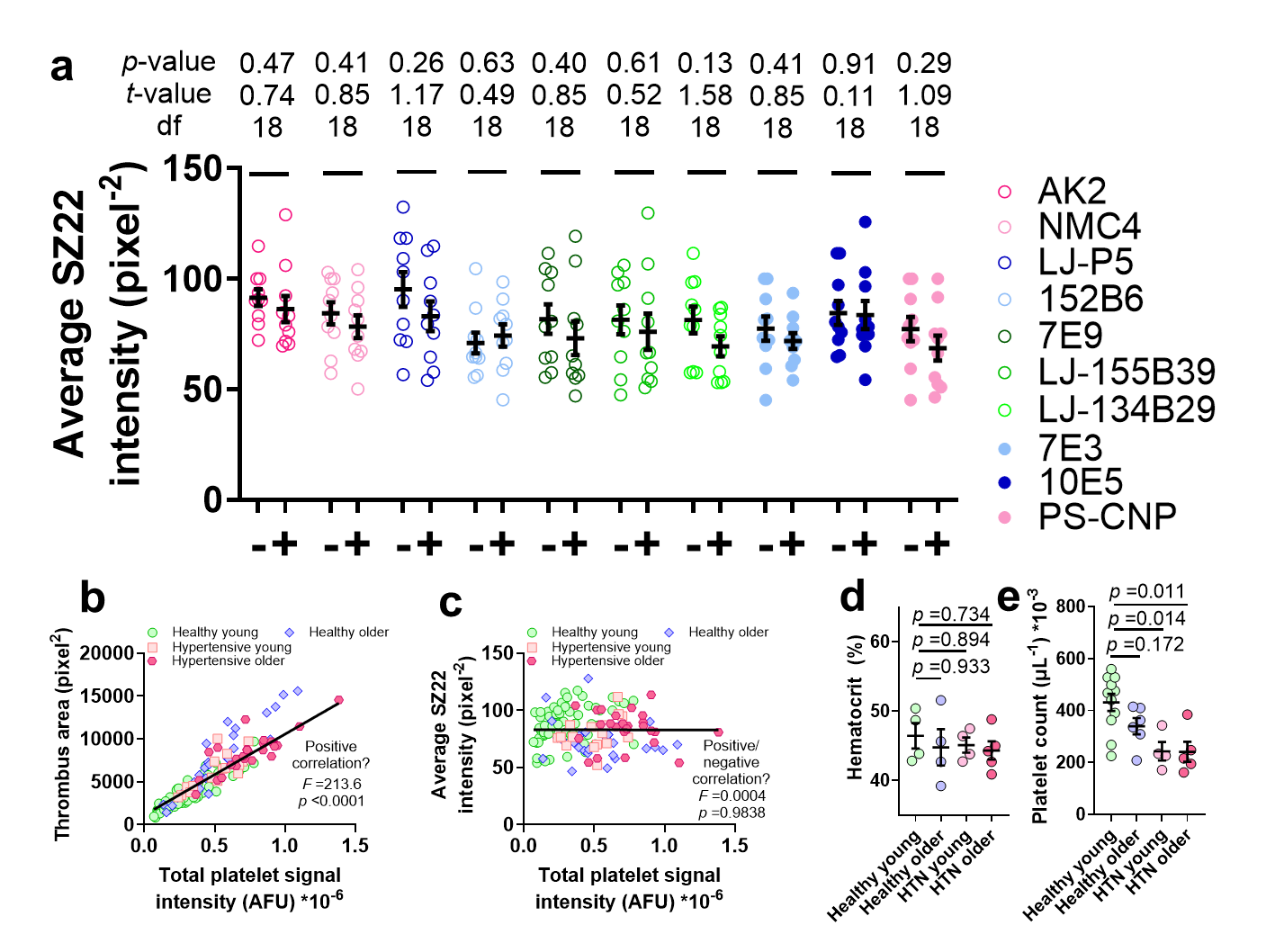

Supplemental Figure 5. Platelet density in the biomechanical thrombi and blood test results. (a) Scatter plots with mean±s.e.m. (n =5; each subject renders two data points acquired from runs using Sensor Set 1 and Set 2, respectively) of the fluorescent signal intensity of SZ22 in the absence and presence of different inhibitors. Student’s t-test was performed for data comparison, with *p*-values, *t*-values and degrees of freedom annotated on the figure. (b,c) Scatter plot of thrombus area (b) and average signal intensity of SZ22 (c) vs. total platelet signal intensity, with blood of healthy young, healthy older, hypertensive young and hypertensive older adult donors (n =33, 14, 9, 13 subjects, respectively; each subject renders two data points acquired from runs using Sensor Set 1 and Set 2, respectively). Solid lines: linear fitting of all data points, with two-sided regression slope test performed to assess whether a positive or negative correlation exists. (d,e) Scatter plots with mean±s.e.m. (n ≥4) of the hematocrit (d) and platelet count (e) of blood from healthy young, healthy older, hypertensive young and hypertensive older adult donors (n =4, 4, 4, 5, respectively). *P*-values: results of one-way ANOVA (F-value =0.258 (d), 6.039 (e); degrees of freedom =16 (d), 25 (e), respectively) and multiple comparison.

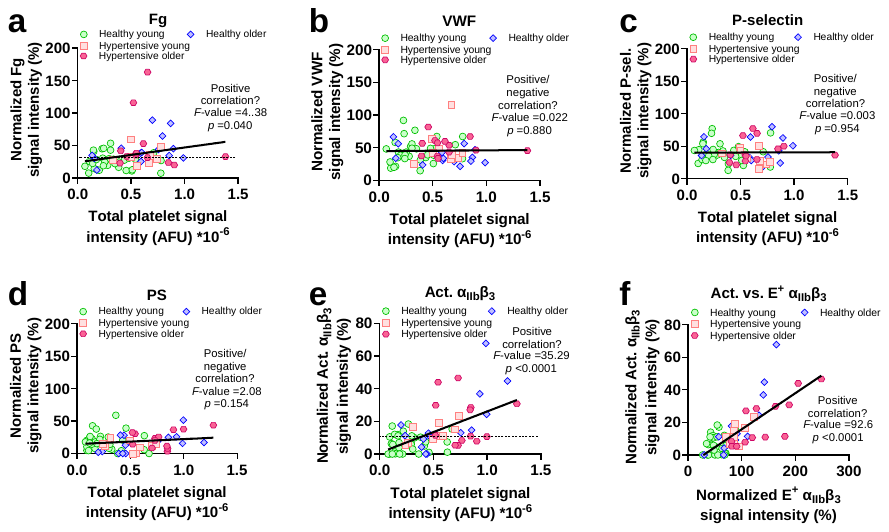

Supplemental Figure 6. Inspecting the correlation between different outputs of the thrombus profile. (a-e) Scatter plots of normalized signal intensity of Fg (a), VWF (b), P-selectin (c), PS (d), Act. α_IIb_β_3_ (e) vs. total platelet signal intensity, using blood from healthy young, healthy older, hypertensive young, and hypertensive older adult donors (n =33, 14, 9, 13, respectively). Dash lines: normalized Fg (a) and Act. α_IIb_β_3_ (e) signal intensity threshold values that best separate healthy young and other groups. (f) Scatter plots of normalized Act. α_IIb_β_3_ signal intensity vs. normalized E^+^ α_IIb_β_3_ signal intensity, with blood of healthy young, healthy older, hypertensive young and hypertensive older adult donors (n =33, 14, 9, 13, respectively). Solid lines: linear fitting of all data points, with two-sided regression slope test performed to assess whether a positive or negative correlation exists.

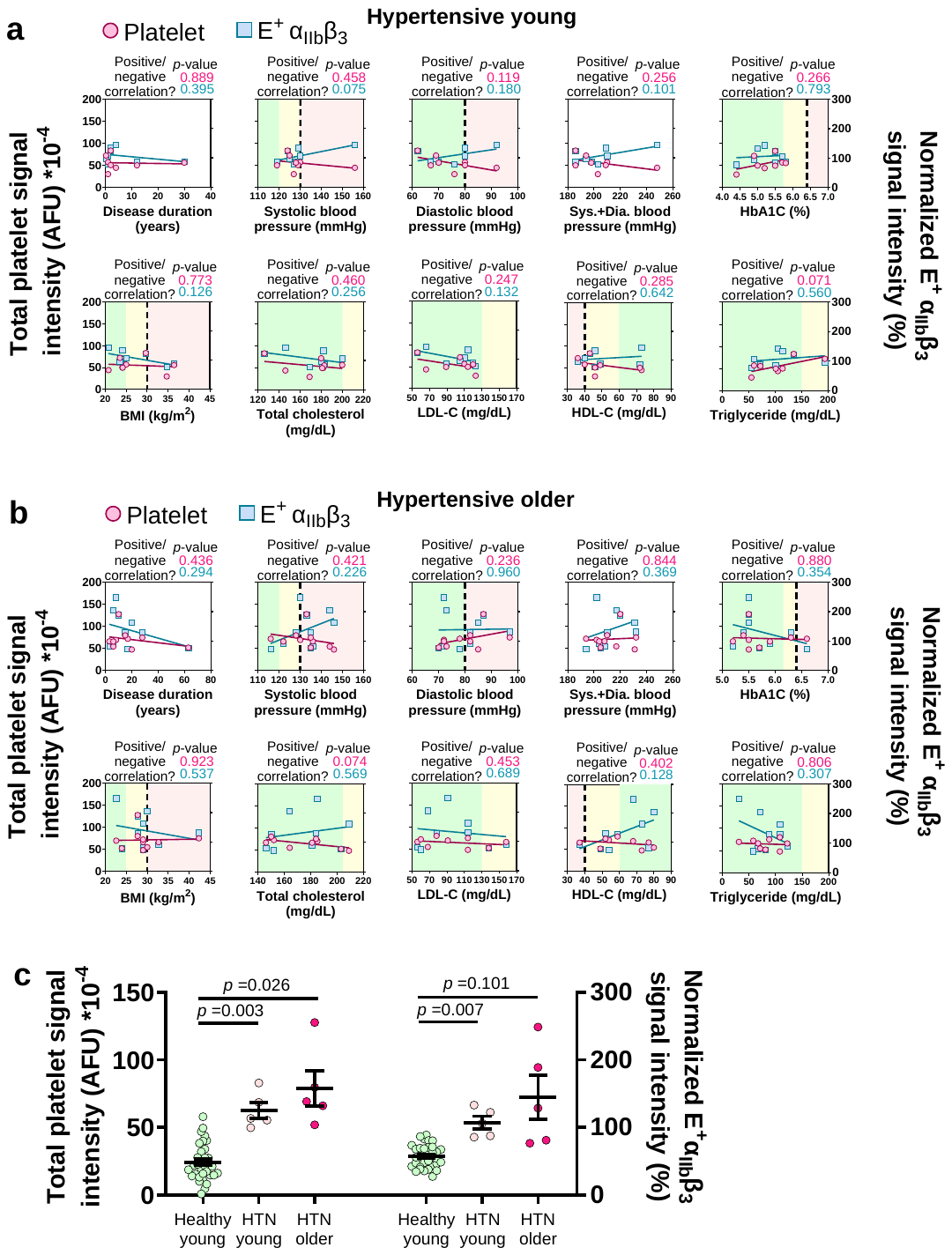

**Supplemental Figure 7. Studying the correlation between thrombus profile abnormality and healthy conditions in hypertension patients.** (**a**,**b**) Scatter plots and linear fits (*P*-values: results of two-sided regression slope test) of total platelet signal intensity and normalized E^+^ α_IIb_β_3_ signal intensity of thrombi vs. disease duration, systolic and diastolic blood pressures and their sum, HbA1C, BMI, total cholesterol, LDL-C, HDL-C and triglyceride levels in young (a) and older (b) hypertension patients. Green, yellow and red background colors indicate normal, borderline abnormal and pathologically abnormal ranges, respectively. (**c**) Comparing the total platelet signal intensity and normalized E^+^ α_IIb_β_3_ signal intensity of thrombi from the blood of healthy young subjects (n =33) and from hypertensive young (n =5) or older (n =5) subjects who have normal systolic and diastolic blood pressures and HbA1C, BMI, total cholesterol, LDL-C, HDL-C and triglyceride levels. *P*-values: results of one-way ANOVA (*left*: F-value =18.4; degrees of freedom =42; *right*: F-value =7.6; degrees of freedom =42) and multiple comparison.

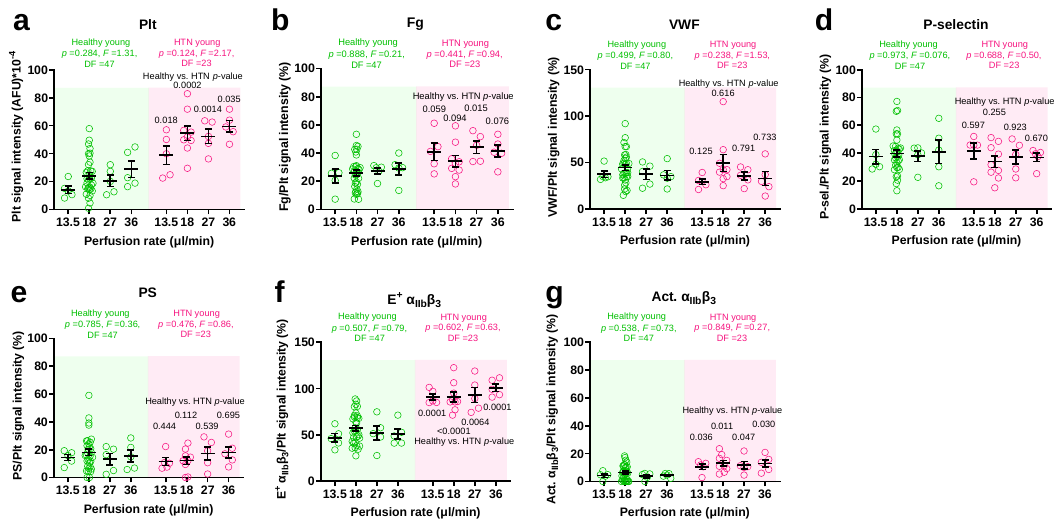

**Supplemental Figure 8. Testing the effect of changing flow perfusion rate on the thrombus profiling results.** The perfusion rate was changed from the original 18 μl/min (data acquired from Fig. 4c; n =33 for healthy young and n =9 for hypertensive (HTN) young) to 13.5, 27 and 36 μl/min, respectively. Then total platelet signal intensity (a) and the normalized signal intensity of Fg (b), VWF (c), P-selectin (d), PS (e), E^+^ α_IIb_β_3_ (f) Act. α_IIb_β_3_ (g) were acquired from n =5 healthy young (*left*, *green points*) and n =5 HTN young (*right*, *magenta points*) subjects and presented as scatter plots with mean±s.e.m.. *P*-values annotated on the graphs are results of two-sided t-tests comparing the result of each HTN young group with that of the healthy young group with the identical perfusion rate. One-way ANOVA was performed to compare the results acquired under different perfusion rates, with the outcome annotated on the graphs (green for healthy young, magenta for HTN young).

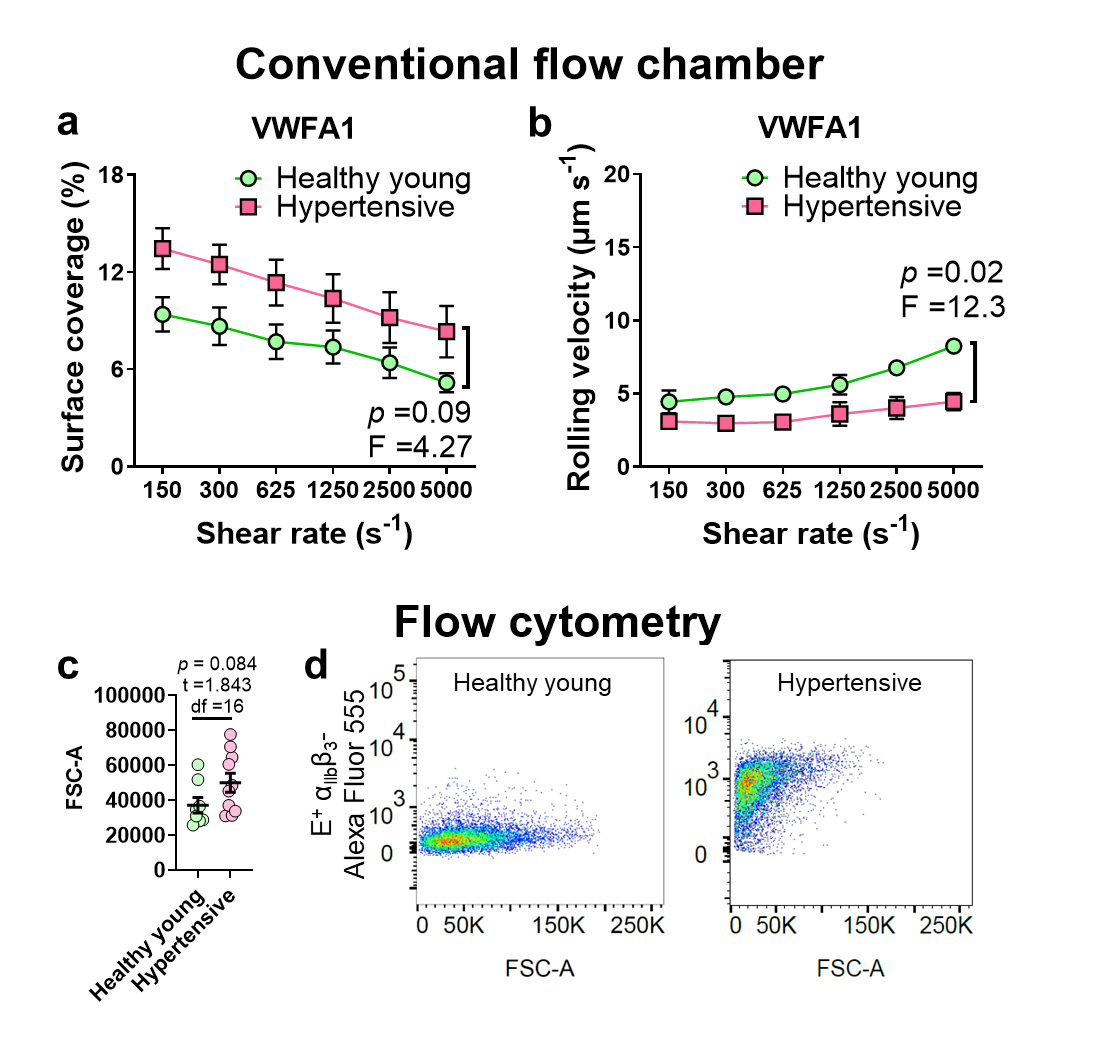

Supplemental Figure 9. Extra conventional flow chamber and flow cytometry data on hypertension patients’ platelets. (a,b) Mean±s.e.m. (n =3) of surface coverage (a) and rolling velocity (b) vs. shear rate of platelets perfused over a surface pre-coated with 30 μg mL^-1^ VWFA1 solution for 1 h. *P*- and *F*-values are results of two-way ANOVA. (c) Scatter plots with mean±s.e.m. of the size (FSC-A, forward scatter area) of healthy young subjects’ (n =8) and hypertension patients’ (n =10) platelets. Statistical comparison was conducted by two-tailed Student’s t-test, with *p*-value, *t*-value and degree of freedom (df) annotated on the figure. (d) Representative scatter plots of E^+^ α_IIb_β_3_ signal intensity (indicated by MBC370.2-Alexa Fluor 555) vs. FSC-A of a healthy young subject’s (*left*) and a hypertension patient’s (*right*) platelets. A total of 10,000 platelets were tested in each panel.

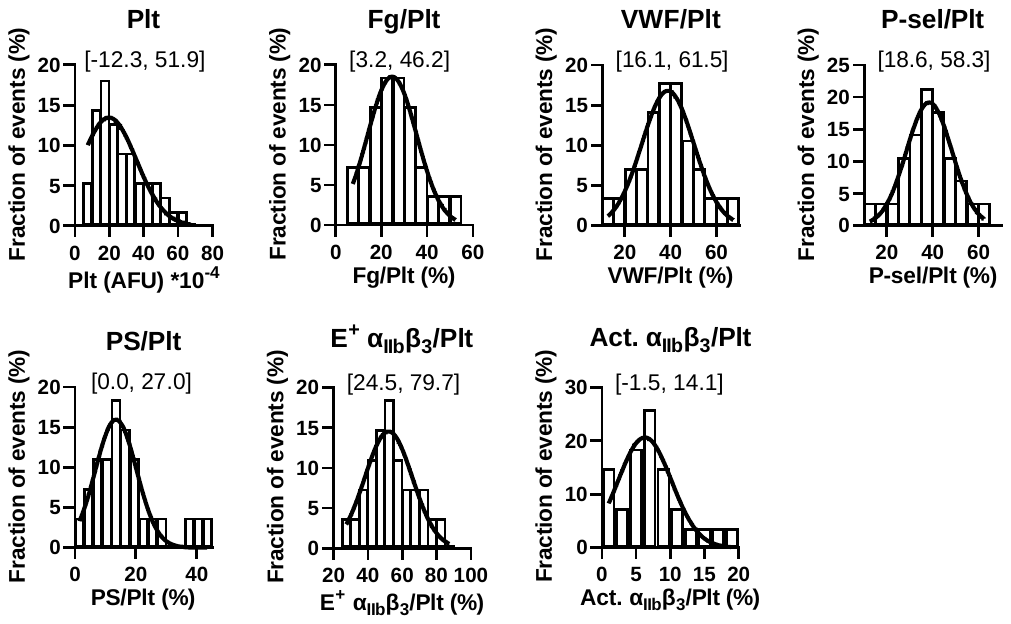
 Supplemental Figure 10. Histograms (bars) of the values of the healthy young adult subjects’ thrombus profiles and the Gaussian fits (curves). The range of mean±2S.D. of each Gaussian fit is annotated.

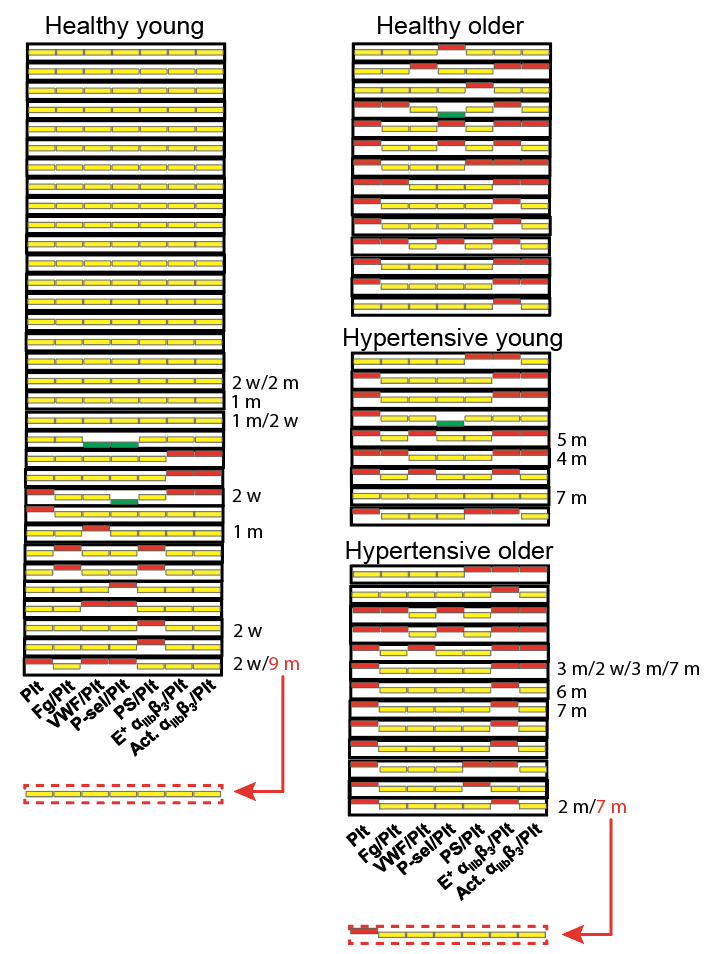

Supplemental Figure 11. Personal thrombus barcodes of healthy young, healthy older, hypertensive young, and hypertensive older adult subjects. For easier visualization, bars indicating ‘abnormally high’, ‘normal’ and ‘abnormally low’ are respectively marked by red, yellow and green. Annotations on the right side of the barcodes indicate that a total of 14 subjects were randomly picked for re-testing after the indicated interval time (‘w’ = week; ‘m’ = month). Among the 21 repeated tests, two (marked in red) rendered different thrombus barcodes, which were shown at the bottom.

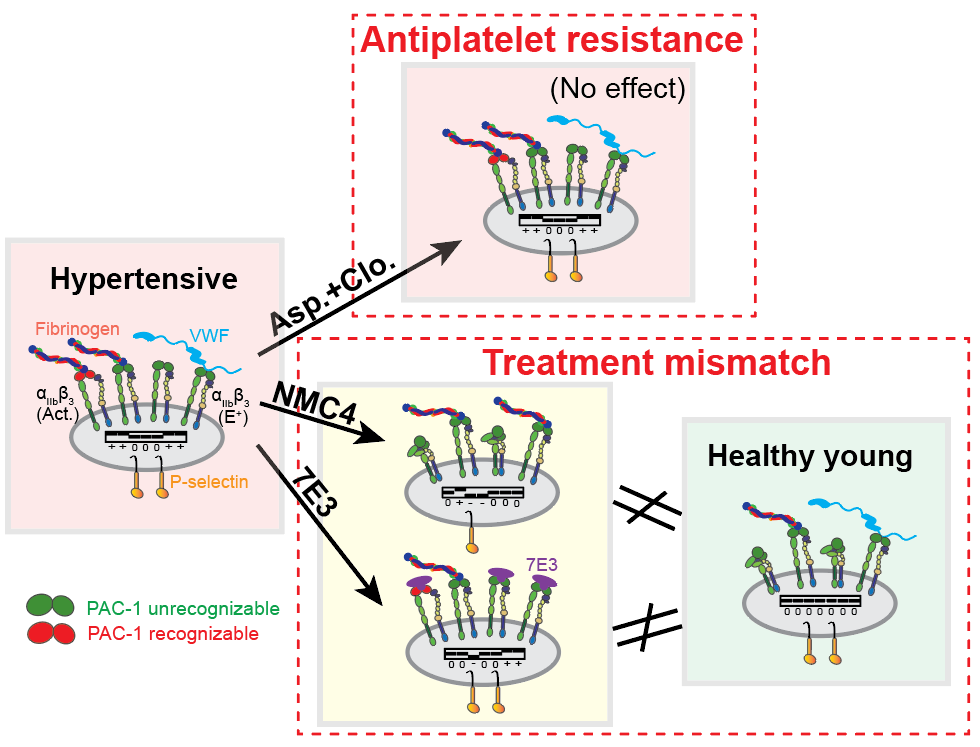

Supplemental Figure 12. Studying the biomechanical thrombogenesis of hypertensive patients reveals a new mechanism of antiplatelet resistance and a ‘treatment mismatch’. The status of platelets in the biomechanical thrombi generated by hypertensive patients’ blood with different treatments is compared with that of platelets in the biomechanical thrombi of healthy young adults’ blood. The thrombus barcodes under different conditions are illustrated inside each platelet. The relative expression level and enrichment of molecules are qualitatively reflected by their numbers in the figure. The engagement of 7E3 at the head of integrin α_IIb_β_3_ blocks the integrin for ligand binding. Aspirin and clopidogrel in combination have no effect on biomechanical thrombogenesis, indicating a new mechanism of antiplatelet resistance. After NMC4 or 7E3 treatment, the status of hypertensive patients’ platelets in the biomechanical thrombus does not match that of healthy young subjects’ platelets, showing a treatment mismatch denoted by a ‘≠’ sign. The head of integrin α_IIb_β_3_ colored green or red respectively denotes the integrin being unrecognizable or recognizable by PAC-1.

**Supplementary Tables**

| **Inhibitor** | AK2 | NMC4 | LJ-P5 | 152B6 | 7E9 | LJ-155B39 | LJ-134B29 | 7E3 | 10E5 |
| --- | --- | --- | --- | --- | --- | --- | --- | --- | --- |
| **IC50 (µg/mL)** | 0.008±  0.002 | 0.266±  0.037 | 0.0086±  0.0018 | 0.0031±  0.0006 | 0.699±  0.182 | 10.37±  8.31 | 13.69±  3.50 | 0.0011±  0.0004 | 0.0011±  0.0006 |
| **R (%)** | 0 | 0 | 4.52±  1.63 | 13.81±  2.58 | 4.98±  2.02 | 20.24±  8.51 | 41.80±  4.51 | 0 | 0 |
| **HillSlope** | -1.76±  0.49 | -1.37±  0.18 | -1.07±  0.27 | -0.920±  0.142 | -0.754  ±  0.123 | -0.978  ±  0.655 | -1.93±  0.89 | -0.618  ±  0.173 | -0.472  ±  0.102 |

Supplemental Table 1. Parameters derived from fitting the thrombus residue size vs. concentration data of different mAbs to the Hill equation.

| **Readout** | | Plt | Fg/Plt | VWF/Plt | P-sel/Plt | PS/Plt | E^+^ α_IIb_β_3_/Plt | Act. α_IIb_β_3_/Plt |
| --- | --- | --- | --- | --- | --- | --- | --- | --- |
| ***Versus* Plt** | **Slope**  **(%*10^-6^)** | N.A. | 25.4±  11.53 | 6.5±  8.5 | -1.3±  7.8 | 8.9±  5.7 | 106.6±  13.1 | 22.5±  3.9 |
|  | **Slope non-zero *p*-value** | N.A. | 0.040 | 0.880 | 0.954 | 0.154 | <0.0001 | <0.0001 |
|  | **Spearman rank correlation**  ***p*-value** | N.A. | 0.007 | 0.20 | 0.77 | 0.18 | <0.0001 | <0.0001 |
|  | **Kendall’s tau correlation**  ***p*-value** | N.A. | 0.007 | 0.18 | 0.82 | 0.20 | <0.0001 | <0.0001 |
| **Optimal separation threshold** | | 3.85*10^5^ | 31.1% | N.A. | N.A. | N.A. | 75.0% | 10.6% |
| **Sensitivity-specificity** | | (83%, 82%) | (69%, 79%) | N.A. | N.A. | N.A. | (86%, 85%) | (69%, 76%) |

Supplemental Table 2. Statistical analysis on the thrombus profiling results to identify potential biomarkers of biomechanical thrombosis. In ‘sensitivity-specificity’, the two percentage numbers respectively represent the specificity and sensitivity of the metrics in separating the healthy young group and the hypertensive and/or older age groups.

|  | Age | Gender | Race | Hispanic/Latino | Hypertension type | Disease duration (years) | Treatment duration (years) | SBP (mmHg) | DBP (mmHg) | Medication (daily) | BMI (kg/m^2^) | HbA1C (%) | TC (mg/dL) | LDL-C (mg/dL) | HDL-C (mg/dL) | TGs (mg/dL) |
| --- | --- | --- | --- | --- | --- | --- | --- | --- | --- | --- | --- | --- | --- | --- | --- | --- |
|  | 43 | F | White | No | Primary | 30 | 30 | 128 | 70 | Losartan 100 mg, hydrochlorothiazide, 25 mg | 36.5 | 5.8 | 183 | 120 | 49 | 72 |
|  | 68 | F | White | No | Primary | 20 | 20 | 146 | 85 | Enalapril 10 mg | 29 | 5.5 | 209 | 114 | 73 | 108 |
|  | 60 | F | White | No | Primary | 8 | 8 | 130 | 72 | Lisinopril 40 mg, Amlodipine 5 mg | 22.6 | 5.5 | 185 | 91 | 88 | 31 |
|  | 46 | F | Other | Yes | Primary | 12 | 5 | 119 | 67 | Amlodipine 5 mg, Hydrochlorothiazide 25 mg | 24.2 | 5.5 | 181 | 89 | 72 | 101 |
|  | 53 | F | Black | No | Primary | 28 | 25 | 135 | 97 | Metoprolol succinate XL 50 mg, Amlodipine 5 mg | 42.3 | 6.3 | 184 | 114 | 52 | 88 |
|  | 39 | F | Black | No | Primary | 2 | 2 | 124 | 62 | Losartan 100 mg, Amlodipine 5 mg | 29.7 | 5.5 | 126 | 56 | 43 | 135 |
|  | 77 | M | White | No | Primary |  |  |  |  | Lisinopril 40 mg, Amlodipine 5 mg daily |  |  |  |  |  |  |
|  | 55 | F | White | No | Primary | 3 | 3 | 136 | 72 | Losartan 50 mg, Spironolactone 50 mg | 29 | 5.2 | 146 | 56 | 77 | 67 |
|  | 64 | F | Black | No | Primary | 6 | 6 | 144 | 73 | Hydrochlorothiazide 25 mg, Minoxidil 2.5 mg | 30 |  | 164 | 68 | 82 | 71 |
|  | 44 | M | White | No | Primary | 1 | 1 | 130 | 80 | Metaprolol succinate 25 mg | 25 | 5.7 | 200 | 110 | 40 | 60 |
|  | 80 | M | Black | No | Primary | 63 | 19 | 135 | 70 | Enalapril 10 mg, Amlodipine 10 mg, Hydrochlorothiazide 25 mg | 23.9 | 5.7 | 203 | 138 | 49 | 81 |
|  | 43 | M | Asian | No | Primary | 4 | 4 | 156 | 92 | Amlodipine 10 mg | 20.8 | 5.2 | 146 | 66 | 73 | 105 |
|  | 52 | M | Asian | No | Primary |  |  |  |  | Losartan 100 mg |  |  |  |  |  |  |
|  | 56 | F | White | No | Primary | 15 | 15 | 128 | 82 | Valsartan 80 mg | 27.7 | 5.4 | 150 | 78 | 59 | 108 |
|  | 34 | M | White | No | Primary | 1 | 1 | 127 | 76 | Lisinopril 10 mg | 34.7 | 4.4 | 169 | 123 | 46 | 55 |
|  | 63 | F | White | No | Primary | 17 | 17 | 116 | 78 | Carvedilol 6.25 mg twice, Enalapril 10 mg | 29 | 6.6 | 152 | 60 | 54 | 58 |
|  | 51 | F | Other | Yes | Primary | 10 | 6 | 133 | 87 | Losartan 100 mg | 27.7 | 5.5 |  |  |  |  |
|  | 23 | M | White | No | Primary | 2 | 1 | 129 | 80 | Amlodipine 5 mg | 24.1 | 5 | 182 | 114 | 46 | 114 |
|  | 32 | M | Asian | No | Primary | 0.25 | 0.25 | 125 | 69 | Losartan 50 mg | 23.5 | 4.9 | 180 | 105 | 36 | 194 |
|  | 51 | F | White | No | Primary |  |  |  |  | Losartan 100 mg |  |  |  |  |  |  |
| 21 | | F | White | No | Primary |  |  |  |  | Prazosin 10 mg |  |  |  |  |  |  |
| 52 | | M | Asian | No | Primary | 6 | 6 | 122 | 82 | Losartan 50 mg | 32.7 | 5.9 | 181 | 158 | 37 | 122 |

Supplemental Table 3. Demographic and health information of the hypertension patients enrolled in this study. Abbreviations: SBP: systolic blood pressure; DBP: diastolic blood pressure; HbA1C: hemoglobin A1c; TC: total cholesterol; HDL-C: high-density lipoprotein cholesterol; LDL-C: low-density lipoprotein cholesterol; TGs: triglycerides. The cells marked in red indicate poorly controlled blood pressure (high SBP and/or DBP), obesity (high BMI) or Type 2 diabetes (high HbA1C).
